# Selecting short DNA fragments in plasma improves detection of circulating tumour DNA

**DOI:** 10.1101/134437

**Authors:** Florent Mouliere, Anna M. Piskorz, Dineika Chandrananda, Elizabeth Moore, James Morris, Christopher G. Smith, Teodora Goranova, Katrin Heider, Richard Mair, Anna Supernat, Ioannis Gounaris, Susana Ros, Jonathan C. M. Wan, Mercedes Jimenez-Linan, Davina Gale, Kevin Brindle, Charles E. Massie, Christine A. Parkinson, James D. Brenton, Nitzan Rosenfeld

## Abstract

Non-invasive analysis of cancer genomes using cell-free circulating tumour DNA (ctDNA) is being widely implemented for clinical indications. The sensitivity for detecting the presence of ctDNA and genomic changes in ctDNA is limited by its low concentration compared to cell-free DNA of non-tumour origin. We studied the feasibility for enrichment of ctDNA by size selection, in plasma samples collected before and during chemotherapy treatment in 13 patients with recurrent high-grade serous ovarian cancer. We evaluated the effects using targeted and whole genome sequencing. Selecting DNA fragments between 90-150 bp before analysis yielded enrichment of mutated DNA fraction of up to 11-fold. This allowed identification of adverse copy number alterations, including *MYC* amplification, otherwise not observed. Size selection allows detection of tumour alterations masked by non-tumour DNA in plasma and could help overcome sensitivity limitations of liquid biopsy for applications in early diagnosis, detection of minimal residual disease, and genomic profiling.

## Text

Analysis of circulating tumour DNA (ctDNA) by non-invasive sampling of cell-free DNA from plasma is now becoming an important tool in oncology for molecular stratification, monitoring tumour burden and analysis of genomic evolution during treatment^1–3^. Analysis of ctDNA is technically challenging, as ctDNA is often present in low concentrations in plasma and is mixed with cell-free DNA (cfDNA) of non-cancerous origin, which is generally present at much higher concentrations. In patients with advanced cancers, the median concentration of ctDNA can reach 10% or more of the total cfDNA, but this fraction is much lower in earlier stage cancer, and ctDNA may rapidly decrease following initiation of systemic treatment or surgery^1,2,4,5^. Various strategies have been proposed to improve the sensitivity of ctDNA analysis^2^. Such methods generally focus on a small subset of the genome, such as hot-spot PCR-based assays or ultra-deep sequencing across gene panels^2,6–9^. Recent observations that ctDNA fragments may be shorter than non-tumour cfDNA in plasma has led to suggestions that these differences may be exploited to enrich for the tumour-specific signal in plasma DNA^10–14^. *In-silico* analysis of ctDNA size differences has been used to enhance the signal for chromosomal changes^13^. However, physical size selection to filter out non-tumour DNA prior to large-scale genomic sequencing has not been demonstrated. Therefore, we tested the hypothesis that selecting DNA fragments of specific sizes could improve the sensitivity of detecting genomic alterations in cfDNA from plasma of cancer patients, enabling the detection of point mutations and copy number alterations that are previously undetectable^11^.

Previous reports using paired-end sequencing reads revealed that cell-free DNA is mostly distributed around a mode at 167 bp, a length that could correspond to the chromatosome (core histones + linker)^15,16^. This size distribution pattern is characteristic of a caspase-dependant cleavage, therefore it was hypothesized that apoptosis releases a large fraction of cfDNA into the bloodstream ^14,15,17,^. Previous studies in non-invasive prenatal testing have explored the potential of size selection for enriching the fetal DNA fraction in maternal plasma with both physical and *in-silico* methods^14,18–20^. However, this analysis of the size distribution cannot be easily generalised to tumour-derived fragments as the characterisation of ctDNA-specific patterns requires analysing fragment sizes of DNA with tumour-derived alterations^13^. Plasma samples from xenografted animal models have shown ctDNA to be highly fragmented below 167 bp^10^ and this distribution with a mode at 145 bp has been then confirmed with PCR, atomic force microscopy and recently with whole-genome sequencing^12,21^. If the distribution differs between tumour-derived and non-tumour derived fragments, sieving fragments by their size could reduce the (often overwhelming) fraction of non-tumour cfDNA and improve the signal to noise ratio in downstream analysis.

We first analysed paired-end reads from shallow whole genome sequencing (sWGS) of plasma DNA from animal models xenografted with a human ovarian cancer cells, and confirmed that the distribution of ctDNA differed in this model from non-tumour cfDNA (Fig. 1a, b). ctDNA in this model was enriched in the size range between 90 and 150 bp, while non-tumour cfDNA was dominant at sizes greater than 150 bp, and peaked at ∼166 bp, similar to previous observations in animal models and in human samples ^12,15,21,22^. Based on these observations we used an automated electrophoresis agarose gel selection method to isolate DNA fragments between 90 and 150 bp for downstream analysis. We sequenced size-selected DNA using both sWGS and tagged-amplicon deep sequencing (TAm-Seq^23^), in 26 samples collected from 13 patients with high-grade serous ovarian cancer (HGSOC), and compared the results to those of the same samples without size selection. For each patient, two plasma samples were collected: one pre-treatment, when levels of ctDNA were generally high, and another several weeks after the start of treatment, when levels of ctDNA were often much lower due to treatment^24^. Analysis of the distribution of fragment lengths after *in-vitro* size selection and sWGS indicated that 96% of resulting reads were in the selected range (Fig. 1c). For two of the patients, the distribution of fragment sizes obtained by sWGS without size-selection exhibited a degraded pattern of cfDNA, without a prominent peak at 166 bp or the 10-bp increment peaks, which has been observed previously^12,13,15^, and is present in all the other cases in this study (Suppl. Fig. 1).

**Figure 1:**
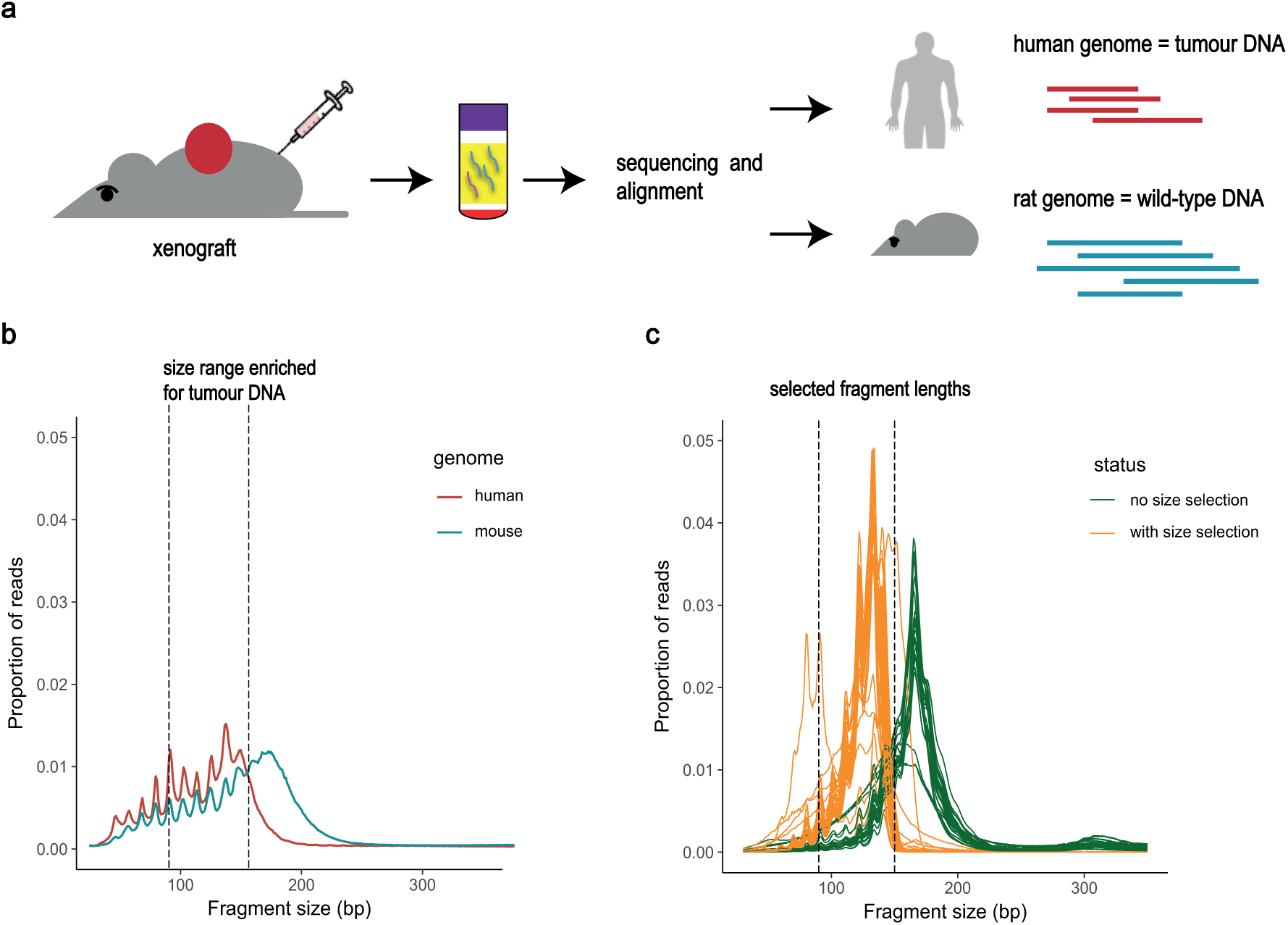
Plasma DNA originating from tumour and non-tumour cells have different sizes, enabling specific enrichment for ctDNA. a. Using an animal model with xenografted cells enabled the discrimination of DNA fragments released by the cancer cells (corresponding to the human DNA) from the DNA fragments released by the wild-type cells (corresponding to the rat DNA). b. Size distribution, assessed by sWGS, of DNA fragments from a plasma sample of a rat xenografted with a human glioblastoma tumour. c. Size distribution of DNA fragments from 26 plasma samples included in this study, assessed by sWGS. In green are the DNA fragments of the samples without size-selection, and in orange after size-selection. The two dotted vertical lines indicate the size selection range between 90 bp and 150 bp.

Analysis of somatic copy number aberrations (SCNAs) was carried out with sWGS on all plasma samples before and after size selection of DNA fragments between 90 and 150 bp (Fig. 2a). One case is exemplified in Fig. 2b and 2c (see further data in Supplementary Fig. 2). Without size selection, a small number of SCNAs were detected with sWGS in DNA isolated from plasma collected 3 weeks after initiation of treatment from patient OV04-83 (Fig. 2b). These included focal amplifications in chromosomes 8p, 14p, 17, and 19q that were observed in this sample at very low levels (Suppl. Table 2). Analysis of the same DNA sample following size-selection for short DNA fragments revealed an increase in the level of these detected SCNAs, in addition to multiples other SCNAs that were not observed without size selection (Fig. 2c). The same pattern of SCNAs and focal amplifications was observed in DNA from plasma collected from the same patient before initiation of the treatment, when the fraction of tumour DNA in the plasma was higher (Suppl. Fig. 2).

**Figure 2:**
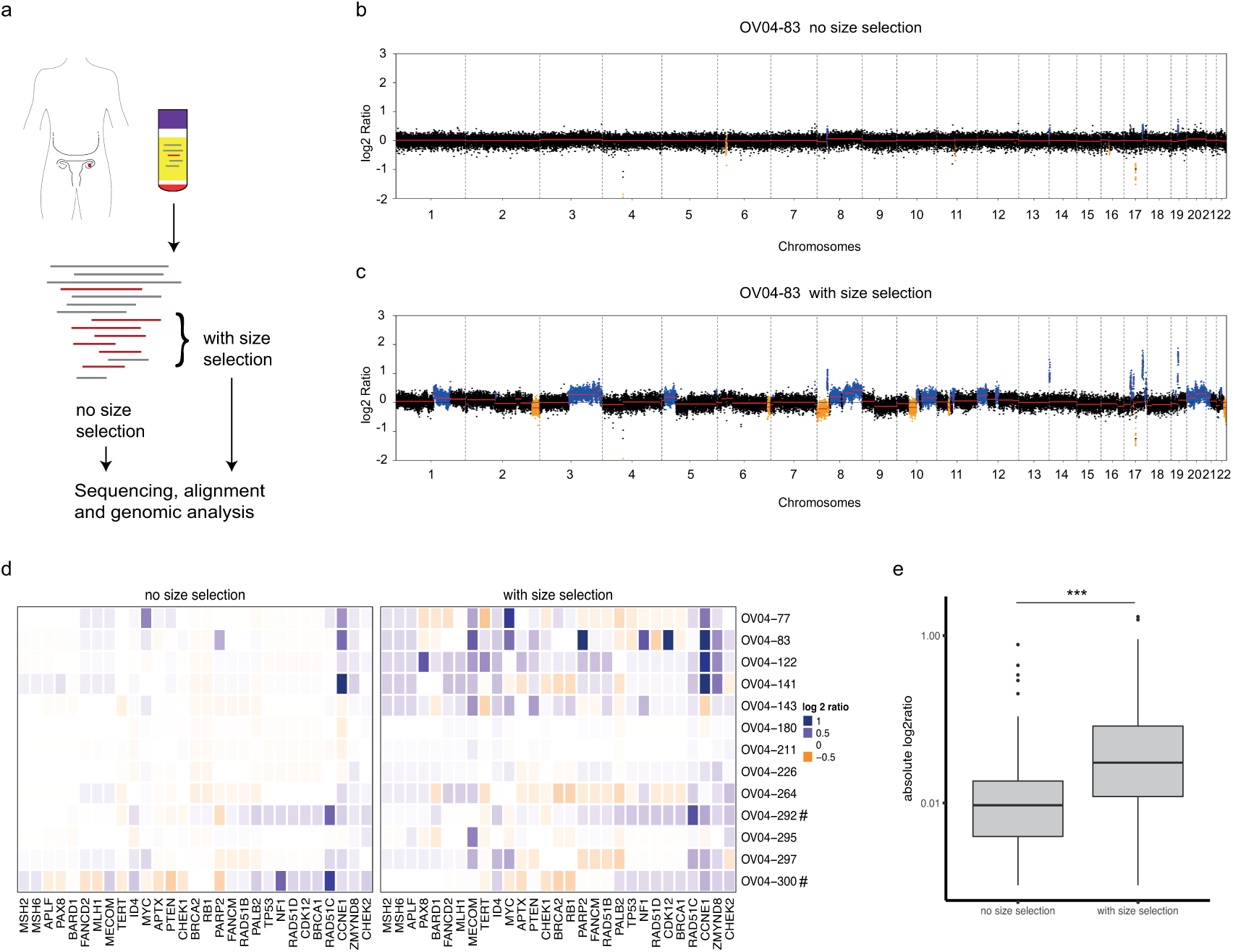
Recovery of short cfDNA fragments enriches for the representation of the cancer genome. a. Samples collected from 13 patients with HGSOC were analysed either with or without filtering by size selection. b. SCNA analysis based on the log2 ratios of regions along the genome of DNA extracted from a plasma sample collected during treatment for patient 0V04-83. c. The same analysis of the same sample with size selection of fragments between 90 bp and 150 bp. Inferred amplifications are shown in blue and deletions in orange. d. SCNA analysis of the segmental log2ratio across a list of 29 genes frequently mutated in recurrent ovarian cancer, measured in plasma samples collected during treatment for all 13 patients, without size selection (left) and with size selection (right). The two samples which exhibited a degraded pattern of cfDNA fragmentation were 0V04-292 and 0V04-300 (both labelled by “#”). e. A comparison of the absolute level of log2ratio across the 29 genes of interest indicated a significant difference between the same samples without and with size selection (p = 7.72.10^-9^). The 2 samples with the degraded pattern of cfDNA fragmentation have been excluded from this analysis.

The relative copy number signals in plasma DNA, across a list of 29 genes frequently mutated in HGSOC, were compared with and without size selection of DNA from plasma samples collected during treatment across the cohort of 13 patients. This showed that a large number of SCNAs that were not observed without size selection, could be detected after size selection for shorter DNA fragments, notably as amplifications in key genes such as *NF1, PARP2* and *MYC* (Fig. 2d and Suppl. Fig. 3). More SCNAs could be detected after size selection in 11/13 patients, and the absolute level of the log2ratio was significantly increased after size selection (t-test, p=7.72.10^-9^). The 2 patients for whom the SCNA signal did not increase exhibited a degraded pattern of cfDNA, which could explain why the size selection had not enriched for ctDNA (Fig. 2d and Suppl. Fig. 2).

We next assessed the detection of SCNAs and point mutations, in plasma samples of the 13 patients collected at baseline (before treatment initiation) and 3 weeks after initiation of chemotherapy treatments, with and without size selection of the plasma DNA (Fig. 3a and Suppl. Table 1). The data from the baseline samples, where ctDNA levels are generally higher^24^, was used to identify and confirm genomic changes, which were then studied in the samples after initiation of treatment, with generally, lower levels of ctDNA. The amplitudes of the absolute log2ratio for the SCNAs were higher, and the concordance of the alterations detected between the baseline and post-treatment samples were improved, with size selection of the plasma DNA (Fig. 3b and Suppl. Fig. 3).

**Figure 3:**
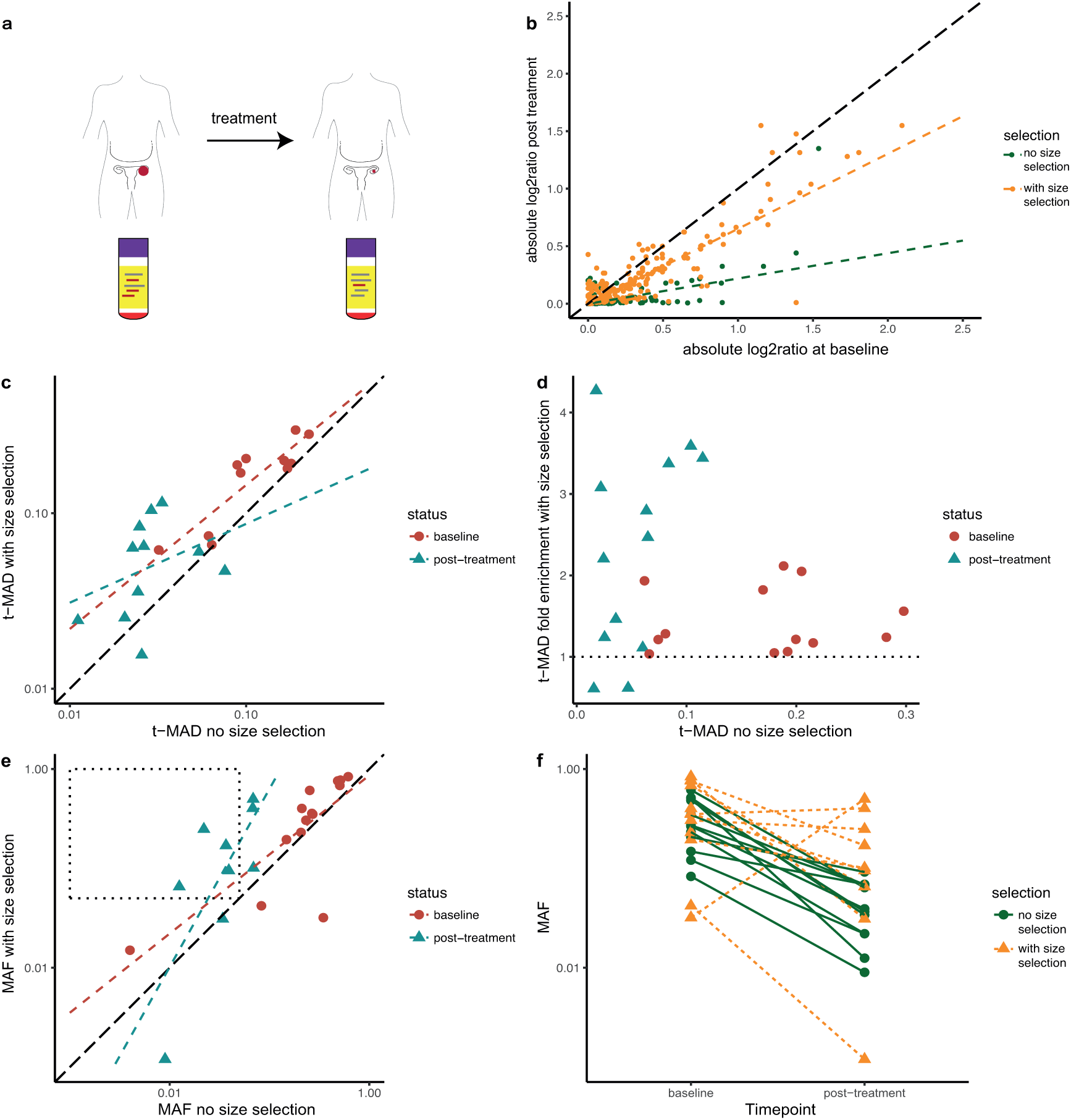
Analysis of the enrichment after size selection in 26 samples sequenced by sWGS and Tagged Amplicon Sequencing (TAm-Seq) revealed relative enrichment in tumour content. a. For each of 13 patients, we compared cfDNA from plasma samples collected before initiation of chemotherapy and 3 weeks or more after initiation of chemotherapy. Each of the 26 plasma samples was analysed with and without size selection. b. Comparison of the absolute value of the segmented log2ratio of the SCNAs called for the plasma samples of patient 0V04-83 collected before and after initiation of the treatment. Data from the samples without size-selection is shown in green, and with size selection in orange. c. The t-MAD score determined from the sWGS with size selection (vertical) was higher than without size selection (horizontal) for most samples, including the samples collected at baseline (red circles) and after initiation of treatment (blue triangles). The 2 samples with no observed enrichment are 0V04-292 and 0V04-300. d. The enrichment factor with size selection, determined by t-MAD, varied per sample but was lower for was samples collected at baseline (red circles), which had high initial t-MAD score, compared to samples collected after treatment (blue triangle). e. The mutant allele fraction (MAF) determined by targeted sequencing with size selection (vertical) was higher than without size selection (horizontal) for most samples, including samples collected at baseline (red circles) and after initiation of treatment (blue triangles). The dotted area highlights samples with low MAF (<5%), where methods such as whole-exome sequencing (at sequencing depth of ∼100x) would not be effective, where size selection enriched the mutant fraction to >5% and therefore accessible for wide-scale analysis. f. Comparison of the MAF detected by TAm-Seq before treatment and after initiation of treatment, as assessed by targeted sequencing, with size selection (yellow triangles) and without size selection (green circles).

Using sWGS data, we converted the amplitude of the CNAs into a quantitative metric called t-MAD (trimmed Mean Absolute Deviation from copy-number neutrality, see *Methods).* Size selection of plasma DNA resulted in a median of 1.5 fold (n=26) increase in the t-MAD score (figure 3c and 3d). However, in the samples collected after the initiation of treatment, when ctDNA content was low, the genome wide enrichment was higher (Fig. 3d), with a median increase of the t-MAD score of 2.9 fold (range: 0.6 - 4.5 fold), except for two samples. For those two samples we observed a decrease in the t-MAD score, and an analysis of the size profile before selection for these samples revealed that the DNA had been heavily fragmented, which could explain why the size selection did not result in enrichment for these cases (Suppl. Fig. 1). In this dataset, we did not identify a differential effect of size selection on the recovery of specific alterations, suggesting that there is a global genome enrichment post size selection. Additionally, size selection notably led to an increased detection of deletions (Suppl. Fig. 2). Analysis with greater sequencing depth, integration of samples from other cancer types and different stages of the disease would help to extend our observations and expand further our understanding of ctDNA biology and fragmentation.

In order to confirm that enrichment for tumour DNA could be observed irrespective of the sequencing approach, we further analysed the mutant allele fractions in the samples using Tagged-Amplicon Sequencing (TAm-Seq). We detected a relative enrichment in the ctDNA fraction in all samples exhibiting a typical pattern of cfDNA fragmentation, with a mode of fragment distribution at 166bp, before size selection (Suppl. Fig. 2). The enrichment effect was below 2-fold in samples collected pre-treatment, when the ctDNA fractions were high in plasma (20%-50% allele fractions for *TP53* mutations as detected by TAm-Seq) (Fig. 3e and 3f). Enrichment of the tumour fraction by size selection was much greater, between 5-fold and 11-fold for most samples, in samples collected approximately 3 weeks after initiation of treatment, when levels of ctDNA (without size selection) were low (ranging from <1% to 5% allele fraction for *TP53* mutations as previously detected by TAm-Seq) (Fig. 3e). For 8 of the 26 plasma samples (31%) we noted a decrease in the allele fractions of mutant *TP53* following size selection; this may be related to loss of rare mutant fragments during the size selection process, or by limitations of this assay for analysis of very short fragments. Increasing the starting amount of material used for size selection, and optimisation of assays for recovery of short fragment, could overcome such limitations.

Methods such as exome-wide sequencing of plasma DNA at multiple time-points of cancer treatment could be effective for the study of cancer evolution and for identification of possible resistance mechanisms to treatment^3,25^. However, analysis with broads-spectrum approaches such as whole-exome sequencing or sWGS are most effective when the ctDNA content is greater than approximately 5%^3,26,27^. In this study, we found that 4 out of 13 cases (31%) where ctDNA levels <5% may have made such analysis uninformative, have been “rescued” by size selection that resulted in >5% mutant allele fractions (Fig. 3e).

These results demonstrate a proof-of-principle that by a simple step of filtering of cfDNA and selection of shorter fragments, it is possible to increase the tumour DNA fraction in plasma cell-free DNA samples. A better understanding of the biology of ctDNA, and their mechanism of release in the circulation, could help select specific fragment ranges depending on their expected origin^28,29^. For example, necrotic cells may release long mutant fragments^14,17^, and thus size selection for fragments of hundreds to thousands of bp may be appropriate for certain samples. Alternatively, a size selection of ultra-short fragments might help for enriching a sample for mitochondrial circulating DNA or bacterial DNA, as their DNA is further fragmented below 100bp^13,30^. Other selection or filtration methods can selectively enrich DNA fragments that are bound to specific molecules for example modified nucleosomes^31^ Determining and accounting for inherent size-biases induced by DNA isolation method that are currently employed for recovery of cfDNA in practice will be important to standardise and optimise methods for liquid biopsy, as large fractions of short or long DNA could be lost at this step. In addition, recent reports have highlighted that different methods of library preparation for sequencing may enable more effective recovery of short DNA fragments from plasma samples, which have led to new observations on cfDNA size distributions^28,30,32^. Such methods should be further investigated to determine if these could help recover more ctDNA.

The size selection process we demonstrated here is based on inherent characteristics of ctDNA in comparison to cfDNA and does not require alteration of these fragments. The enrichment we observed is therefore compatible with any downstream genomic analysis, from locus-specific to wider genomic sequencing. This work shows that sWGS (and by extension, whole exome sequencing) can be performed on plasma DNA samples with low ctDNA content, and that this can facilitate the characterisation of mutations present in plasma at lower allele fractions and with lower sequencing depth. The compatibility of the cfDNA fragment size selection with wide-scale and sensitive genomic analysis could unlock the potential of liquid biopsies for the diagnosis of cancer at an earlier stage, and for the detection of minimal residual disease.

## Methods

### Patients and sample preparation

13 patients were recruited as part of prospective clinical studies at Addenbrooke’s Hospital, Cambridge, UK, approved by the local research ethics committee (REC reference numbers 07/Q0106/63, 08/H0306/61 and 07/Q0106/63). Written informed consent was obtained from all patients and blood samples were collected before and after initiation of treatment with chemotherapeutic agents. DNA was extracted from 2 mL of plasma using the QIAamp circulating nucleic acid kit (Qiagen) according to the manufacturer’s instructions.

### Size selection

Between 8-10 ng of DNA were loaded into a 3% agarose cassette (HTC3010, Sage Bioscience) and size selection was performed on a PippinHT (Sage Bioscience) according to the manufacturer’s protocol.

### TAm-Seq

Tagged-Amplicon Sequencing libraries were prepared as previously described^23^, using primers designed to assess single nucleotide variants (SNV) and small indels across selected hotspots and the entire coding regions of *TP53.* Libraries were sequenced using an HiSeq 4000 (Illumina).

### sWGS

Indexed sequencing libraries were prepared using a commercially available kit (ThruPLEX-Plasma Seq, Rubicon Genomics). Libraries were pooled in equimolar amounts and sequenced on a HiSeq 4000 (Illumina) generating 150-bp paired-end reads. Sequence data was analysed using an in-house pipeline that consists of the following; Paired end sequence reads were aligned to the human reference genome (GRCh37) using BWA-mem following the removal of contaminating adapter sequences^33^. PCR and optical duplicates were marked using MarkDuplicates (Picard Tools) feature and these were excluded from downstream analysis along with reads of low mapping quality and supplementary alignments.

### SCNA analysis

Copy number analysis was performed in R using a modification of the QDNAseq pipeline^34^, as follow: sequence reads were allocated into equally sized (50 kbp) non-overlapping bins throughout the length of the genome. Read counts in each bin were corrected to account for sequence GC content and mappability, and bins overlapping ‘blacklisted’ regions (ENCODE project) were excluded from downstream analysis. After median normalisation of the counts, bins were segmented using both Circular Binary Segmentation and Locus-aware Circular Binary Segmentation algorithms, and an averaged log2R value per bin was calculated. The t-MAD score is calculated as the averaged absolute deviation from log2R = 0 after first trimming bin counts greater than 4 standard deviations from the mean count across all genomic regions.

## Notes

## Acknowledgements

The authors would like to thank all members of the Rosenfeld Lab and Brenton Lab for help and constructive discussion, in particular Francesco Marass, Wendy N. Cooper, Keval Patel, Jenny P.Y. Chan, Mareike Thompson, Lise Barlebo Ahlborn and Irena Hudecovà. The authors would like to also thank the Cancer Research UK Cambridge Institute core facilities for their support, in particular the genomics and biorepository facilities. We would like to acknowledge our patients and our caregivers, and the help and support of the research nurses, trial staff and the staff at Addenbrooke’s Hospital. In particular, we would like to acknowledge Charlotte Hodgkin, Heather Biggs and Karen Hosking. We would like also to acknowledge the support of The University of Cambridge, Cancer Research UK (grant number A11906, A20240) and Hutchison Whampoa Limited. The research leading to these results has received funding from the European Research Council under the European Union’s Seventh Framework Programme (FP/2007-2013) / ERC Grant Agreement n. 337905. This research is also supported by Target Ovarian Cancer and the Medical Research Council through their Joint Clinical Research Training Fellowship for Dr Moore. The funders had no role in study design, data collection and analysis, decision to publish, or preparation of the manuscript.

## Author contributions

FM, AMP, DC, EM, JDB and NR conceptualised and designed the study; FM, AMP, EM, KH, CGS, JCMW, DG, RM, TG, AS, IG, CAP have performed experiments and collected data; DC has conceptualised and designed the t-MAD index and performed sWGS bioinformatics analysis; JM performed TAm-Seq bioinformatics analysis; RM and SR have designed the animal model; MJL performed histopathology revision; FM, AMP, DC, EM, JDB and NR have written the manuscript; all co-authors have critically reviewed the manuscript; FM, AMP, DC, JDB and NR supervised the study.

## Author information

NR, JDB and DG are cofounders, shareholders and officers/consultants of Inivata Ltd, a cancer genomics company that commercialises ctDNA analysis. Inivata Ltd had no role in the conceptualisation, study design, data collection and analysis, decision to publish or preparation of the manuscript. NR and FM are co-inventors of patent applications that describe methods for the analysis of DNA fragments and applications of circulating tumour DNA. Other co-authors have declared no conflict of interests.

## References

1. Siravegna, G., Marsoni, S., Siena, S. & Bardelli, A. Integrating liquid biopsies into the management of cancer. Nat. Rev. Clin. Oncol. (2017). doi:10.1038/nrclinonc.2017.14

2. Wan, J. C. M. et al. Liquid biopsies come of age: towards implementation of circulating tumour DNA. Nat. Rev. Cancer (2017). doi:10.1038/nrc.2017.7

3. Murtaza, M. et al. Non-invasive analysis of acquired resistance to cancer therapy by sequencing of plasma DNA. Nature 497, 108–112 (2013).

4. Bettegowda, C. et al. Detection of Circulating Tumor DNA in Early- and Late-Stage Human Malignancies. Sci. Transl. Med. 6, 224ra24–224ra24 (2014).

5. Dawson, S.-J. et al. Analysis of Circulating Tumor DNA to Monitor Metastatic Breast Cancer. N. Engl. J. Med. 368, 1199–1209 (2013).

6. Diehl, F. et al. Detection and quantification of mutations in the plasma of patients with colorectal tumors. Proc. Natl. Acad. Sci. 102, 16368–16373 (2005).

7. Taly, V. et al. Multiplex Picodroplet Digital PCR to Detect KRAS Mutations in Circulating DNA from the Plasma of Colorectal Cancer Patients. Clin. Chem. 59, 1722–1731 (2013).

8. Newman, A. M. et al. Integrated digital error suppression for improved detection of circulating tumor DNA. Nat. Biotechnol. 34, 547–555 (2016).

9. Khodakov, D., Wang, C. & Zhang, D. Y. Diagnostics based on nucleic acid sequence variant profiling: PCR, hybridization, and NGS approaches. Adv. Drug Deliv. Rev. 105, 3–19 (2016).

10. Mouliere, F. et al. High Fragmentation Characterizes Tumour-Derived Circulating DNA. PLoS One 6, e23418 (2011).

11. Mouliere, F. & Rosenfeld, N. Circulating tumor-derived DNA is shorter than somatic DNA in plasma. Proc. Natl. Acad. Sci. 112, 3178–3179 (2015).

12. Underhill, H. R. et al. Fragment Length of Circulating Tumor DNA. PLOS Genet. 12, e1006162 (2016).

13. Jiang, P. et al. Lengthening and shortening of plasma DNA in hepatocellular carcinoma patients. Proc. Natl. Acad. Sci. 112, E1317–E1325 (2015).

14. Jiang, P. & Lo, Y. M. D. The Long and Short of Circulating Cell-Free DNA and the Ins and Outs of Molecular Diagnostics. Trends Genet. 32, 360–371 (2016).

15. Lo, Y. M. D. et al. Maternal Plasma DNA Sequencing Reveals the Genome-Wide Genetic and Mutational Profile of the Fetus. Sci. Transl. Med. 2, 61ra91–61ra91 (2010).

16. Chandrananda, D. et al. High-resolution characterization of sequence signatures due to nonrandom cleavage of cell-free DNA. BMC Med. Genomics 8, 29 (2015).

17. Jahr, S. et al. DNA fragments in the blood plasma of cancer patients: quantitations and evidence for their origin from apoptotic and necrotic cells. Cancer Res. 61, 1659–65 (2001).

18. Yu, S. C. Y. et al. Size-based molecular diagnostics using plasma DNA for noninvasive prenatal testing. Proc. Natl. Acad. Sci. U. S. A. 111, 8583–8 (2014).

19. Lun, F. M. F. et al. Noninvasive prenatal diagnosis of monogenic diseases by digital size selection and relative mutation dosage on DNA in maternal plasma. Proc. Natl. Acad. Sci. U. S. A. 105, 19920–5 (2008).

20. Minarik, G. et al. Utilization of Benchtop Next Generation Sequencing Platforms Ion Torrent PGM and MiSeq in Noninvasive Prenatal Testing for Chromosome 21 Trisomy and Testing of Impact of In Silico and Physical Size Selection on Its Analytical Performance. PLoS One 10, e0144811 (2015).

21. Mouliere, F., El Messaoudi, S., Pang, D., Dritschilo, A. & Thierry, A. R. Multi-marker analysis of circulating cell-free DNA toward personalized medicine for colorectal cancer. Mol. Oncol. 8, 927–941 (2014).

22. Thierry, A. R. et al. Origin and quantification of circulating DNA in mice with human colorectal cancer xenografts. Nucleic Acids Res. 38, 6159–6175 (2010).

23. Forshew, T. et al. Noninvasive Identification and Monitoring of Cancer Mutations by Targeted Deep Sequencing of Plasma DNA. Sci. Transl. Med. 4, 136ra68–136ra68 (2012).

24. Parkinson, C. A. et al. Exploratory Analysis of TP53 Mutations in Circulating Tumour DNA as Biomarkers of Treatment Response for Patients with Relapsed High-Grade Serous Ovarian Carcinoma: A Retrospective Study. PLOS Med. 13, e1002198 (2016).

25. Murtaza, M. et al. Multifocal clonal evolution characterized using circulating tumour DNA in a case of metastatic breast cancer. Nat. Commun. 6, 8760 (2015).

26. Heitzer, E. et al. Tumor-associated copy number changes in the circulation of patients with prostate cancer identified through whole-genome sequencing. Genome Med. 5, 30 (2013).

27. Belic, J. et al. Rapid Identification of Plasma DNA Samples with Increased ctDNA Levels by a Modified FAST-SeqS Approach. Clin. Chem. 61, 838–849 (2015).

28. Snyder, M. W. et al. Cell-free DNA Comprises an In Vivo Nucleosome Footprint that Informs Its Tissues-Of-Origin. Cell 164, 57–68 (2016).

29. Ulz, P. et al. Inferring expressed genes by whole-genome sequencing of plasma DNA. Nat. Genet. 48, 1273–1278 (2016).

30. Burnham, P. et al. Single-stranded DNA library preparation uncovers the origin and diversity of ultrashort cell-free DNA in plasma. Sci. Rep. 6, 27859 (2016).

31. Shema, E. et al. Single-molecule decoding of combinatorially modified nucleosomes. Science (80-.). 352, (2016).

32. Vong, J. S. L. et al. Single-Stranded DNA Library Preparation Preferentially Enriches Short Maternal DNA in Maternal Plasma. Clin. Chem. (2017). doi:10.1373/clinchem.2016.268656

33. Li, H. & Durbin, R. Fast and accurate short read alignment with Burrows-Wheeler transform. Bioinformatics 25, 1754–1760 (2009).

34. Scheinin, I. et al. DNA copy number analysis of fresh and formalin-fixed specimens by shallow whole-genome sequencing with identification and exclusion of problematic regions in the genome assembly. Genome Res. 24, 2022–2032 (2014).

